# A genomic foundation model for interpreting human genetic diversity

**DOI:** 10.64898/2026.10.19.760276

**Authors:** QinYi Xu, ZhiKun Wu, Meijing Dong, Tianpeng Bao, Wenyan Nong, Xuanzhao Huang, Hongxi Yang, Yao Zhou, Xianfu Yi, Shijie Zhang, Dandan Huang, Pak Chung Sham, Wenhao Zhou, Mulin Jun Li

## Abstract

Interpreting human genomes requires models that recognize shared biological functions while accounting for variation underlying traits, disease susceptibility and adaptation. Here we present hEVO, a genomic foundation model trained on 4.784 billion distinct haplotype sequences reconstructed from whole genome sequencing (WGS) data for 63,809 individuals selected from 550,237 samples across eight cohorts. Pretraining integrates shared sequence features and naturally occurring allelic differences without clinical or functional labels. Against representatives of the Evo 2, NT, GPN and other genomic model families, hEVO ranked first in 18 of 19 primary comparisons spanning functional annotation, variant interpretation and human evolution. Its pretrained allele scores generalized across all six complementary variant benchmarks without task-specific training, achieving the highest performance across clinical pathogenicity, fine-mapped causal variants and experimentally measured regulatory effects. Retrospective ClinVar reclassification further demonstrated improved identification of pathogenic classifications later downgraded, supporting reassessment of earlier clinical interpretations. Predictive strengths extended to ancestry inference, human accelerated sequences and preference for reported selected alleles. This breadth was achieved with substantially fewer parameters than billion-parameter comparators and low inference overhead in the evaluated workloads. These findings establish human haplotype diversity as a source of transferable biological information for interpreting both the disruption of genome function and the variation retained during human evolution.

## Introduction

The human genome encodes the molecular programmes of development and physiology, while variation within it contributes to differences in traits and disease susceptibility. Interpreting this variation requires understanding how sequence changes affect biological function and why their consequences differ across genetic and environmental contexts. Regulatory changes have altered cellular functions during human evolution (1), and recent selection has reshaped the distribution of trait-associated alleles among populations (2). Some alleles favoured in past environments now contribute to disease risk (3, 4). Molecular function, disease susceptibility and evolutionary fitness are therefore related but non-equivalent aspects of genome interpretation. Even large functional studies measure only part of the relevant sequence and cellular space (5), while uneven population representation leaves much human diversity poorly characterized (6). A central challenge is to learn genomic information that remains useful across these biological questions, including settings for which direct functional evidence is sparse.

Genomic language models learn from different records of biological variation. Evo and Evo 2 (7, 8) use genomes across species to support molecular prediction and sequence design, whereas GPN-MSA (9) and the phylogeny-aware GPN-Star (10) exploit aligned evolutionary histories to estimate functional constraint. Their success demonstrates the value of comparative genomics, but conservation across lineages captures only part of the variation segregating within humans. Nucleotide Transformer (NT) (11) and Genos (12) incorporate human haplotypes or diverse genome assemblies, while MutBERT (13) and UKBioBERT (14) explicitly incorporate population variant information. These approaches establish that human diversity is relevant to sequence learning, leaving open how broadly that diversity must be sampled and how it should inform predictions at variable sites. AlphaGenome (15) addresses a complementary objective, learning from experimental measurements to predict gene expression, splicing and chromatin properties across long genomic contexts. Its molecular outputs offer a different route to variant interpretation from the sequence representations and allele likelihoods learned through genomic language modelling.

The distinction matters because recognizing functional DNA does not guarantee sensitivity to the differences between individual genomes. Models that predict expression across genes can struggle to recover variation between individuals (16), although training on personal genomes can improve this capability (17). Human haplotypes preserve alleles within the sequence backgrounds in which they occur, recording variation shaped by population history, functional constraint and recent adaptation (2, 18). This information is not interchangeable with divergence between species or with isolated pathogenicity labels. We reasoned that learning diverse human sequence backgrounds, then refining predictions at naturally variable sites, could connect functional sequence recognition with allelic interpretation within one model.

Here we present hEVO, a genomic foundation model trained on haplotypes from 63,809 individuals selected from an initial collection of 550,237 whole genome-sequenced participants across eight cohorts. The training resource retains distinct sequence backgrounds while reducing repeated exposure to identical DNA, and a second learning stage refines allele predictions without replacing the sequence representation. We evaluated 34 checkpoints across functional annotation, six clinical and functional variant benchmarks, retrospective clinical reclassification, and human population and evolutionary analyses. With only 87 million or 319 million parameters, hEVO-base or hEVO-large led eligible external models in 18 of 19 primary benchmark comparisons, combining broad predictive utility with modest model size and sequence-only inference. This combination provides a practical foundation for studying human genomic function and variation across questions that extend beyond established pathogenic labels.

## Results

### Human haplotype diversity supports integrated sequence and allelic learning

We assembled a human sequence resource to retain diverse genetic backgrounds while limiting redundant training. The initial collection comprised whole genome sequencing (WGS) data from 550,237 participants from UK Biobank (19), TOPMed (20), the Human Genome Diversity Project (HGDP) (18), the 1000 Genomes Project (1000G) (21), UK10K (22), Singapore 10K (SG10K) (23), the Consortium on Asthma among African ancestry Populations in the Americas (CAAPA) (24) and an internal cohort of healthy Han Chinese individuals. Selection using UK Biobank ancestry codes reduced its European-ancestry dominance to a subset of 25,934 individuals. In 100 kb windows, this subset retained 95.3% of abundance-weighted reference haplotype coverage above a frequency of 0.01% and more than 99.99% above 0.1%, although coverage fell to 77.6% when rarer haplotypes down to 0.001% were included (Fig. 1a). Thus, substantial cohort reduction preserved most frequent local backgrounds while incompletely capturing the rarest diversity. After quality control and duplicate removal, the final resource contained 63,809 individuals, including 21,310 from TOPMed, with eligible samples from the seven non-UK Biobank cohorts retained without representative downsampling (Fig. 1b,c, Supplementary Fig. S1a-c and Supplementary Table S1).

**Figure 1.**
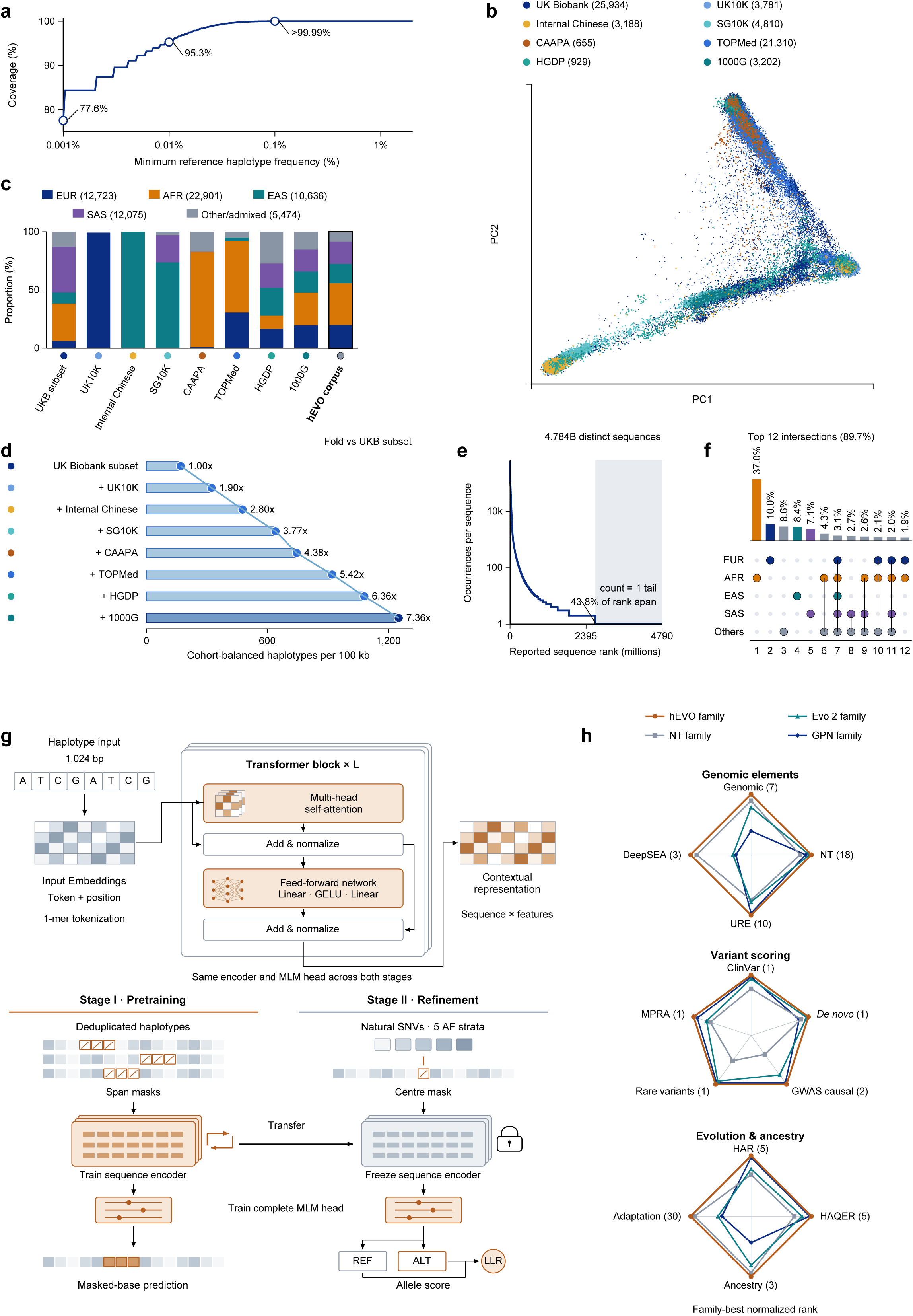
Human haplotype diversity and integrated genomic learning. **a,** Abundance-weighted coverage of local UK Biobank SNV haplotypes by the selected 25,934 individuals across minimum reference haplotype frequencies. Haplotypes were defined from phased SNVs in 100 kb windows, separately from the 1,024 bp training sequences. Coverage is the fraction of eligible haplotype abundance in a 50,000-person reference sample from the full phased UK Biobank cohort that is represented in the selected subset. **b,c,** Genetic principal components (b) and cohort-specific population composition (c) of the final 63,809-individual WGS resource. The final UK Biobank contribution comprised the same 25,934 individuals used in the haplotype coverage analysis. Composition annotations use a calibrated distance-weighted 15-neighbour classifier on PC1-PC6 with 2,791 1000G/HGDP reference individuals. The 3,188 internal Chinese participants are annotated as EAS from cohort information. Accepted AMR, Middle Eastern and Oceanian assignments and unassigned samples are summarized as Others. These categories do not define discrete biological boundaries. **d,** Accumulation of local haplotype diversity as cohorts are added to the selected UK Biobank subset. Each contributing cohort supplies an equal number of chromosome copies within a scenario, so total sample size increases with cohort accumulation. Bars summarize 577 sampled, variant-bearing 100 kb windows, with 95% percentile intervals from 1,000 bootstrap resamples of windows. Displayed fold changes are relative to the selected UK Biobank starting subset. **e,f,** Abundance distribution of 4.784 billion distinct reconstructed sequences (e) and sharing across five ancestry categories (f). Singletons account for 43.8% of distinct sequences. The 12 displayed ancestry intersections account for 89.7%. Sharing is summarized using the original PC1/PC2-based ancestry assignments. **g,** Integrated architecture and two-stage training. Stage I trains the sequence encoder and masked language modelling (MLM) prediction head on distinct haplotype sequences. Stage II freezes the encoder and trains the complete existing head, including its transformation layer, LayerNorm and decoder, using natural SNVs sampled across five frequency strata. Neither stage uses clinical or functional outcome labels. **h,** Family-best profiles across 13 benchmark axes. Within an axis, rank r among N eligible models is transformed to (N − r + 1)/N, with larger radii indicating a better relative rank. Combined axes average the defined component summaries before ranking. Each family contributes its best eligible configuration per axis, so a polygon need not represent one checkpoint. GPN-Star, GPN-MSA and GPN-Promoter belong to the GPN family in this summary.

Integrating the cohorts expanded the diversity available beyond the selected UK Biobank subset. Across 577 sampled, variant-bearing 100 kb windows, mean local haplotype richness increased from approximately 170 to 1,251 as cohorts were added with balanced contributions, a 7.4-fold accumulation (Fig. 1d). Reconstruction and exact deduplication of overlapping 1,024 bp haplotype sequences yielded 4.784 billion distinct strings (Supplementary Fig. S1d,e and Supplementary Table S2). Of these, 43.8% occurred only once, indicating that much of the available sequence diversity would receive little exposure if training followed raw abundance (Fig. 1e). Sequence-sharing annotations further distinguished backgrounds observed in only one ancestry category, comprising 71.1% of distinct strings, from those shared across all five categories, comprising 3.1% (Fig. 1f). Sampling distinct strings equally made both recurring and sparsely observed backgrounds available for learning, rather than allowing identical copies to determine their relative representation.

hEVO uses a nucleotide-level bidirectional Transformer to learn sequence context and predict alleles through a common representation. Stage I learns broad sequence features by reconstructing masked spans of haplotype DNA. Stage II then focuses on naturally occurring single-nucleotide variants (SNVs), sampled across five frequency strata, while preserving the sequence encoder learned in Stage I. Only the complete existing masked language modelling (MLM) prediction head is updated, including its transformation layer, LayerNorm and nucleotide decoder. Neither stage uses clinical pathogenicity or functional labels (Fig. 1g and Supplementary Fig. S2). We trained an 87-million-parameter model, hEVO-base, and a 319-million-parameter model, hEVO-large, to examine the predictive breadth attainable at two compact scales. Each supplies sequence representations for functional analysis and allele probabilities for variant scoring within the same architecture (Supplementary Tables S3 and S30).

The evaluation connected three broad uses of genomic prediction. Functional assessment comprised eight comparisons: two standard sequence benchmark collections, three annotation-based representation analyses, five-class regulatory recognition, unmarked regulatory element (URE) versus enhancer discrimination and multitarget chromatin prediction. Six variant benchmarks tested clinical, association and experimental evidence without fitting classifiers to their labels. Five population and evolutionary comparisons assessed ancestry classification, ancestry-proportion prediction, recognition of two classes of human accelerated sequence and preference for reported selected alleles. hEVO-base or hEVO-large led eligible external models on the primary summary metric in 18 of these 19 comparisons; GENCODE annotation separation was the exception (Figs. 2-4). Three retrospective clinical reclassification outcomes provided an additional longitudinal assessment. The family profiles summarize related comparisons on 13 axes, selecting the best eligible configuration per family and axis (Fig. 1h).

**Figure 2.**
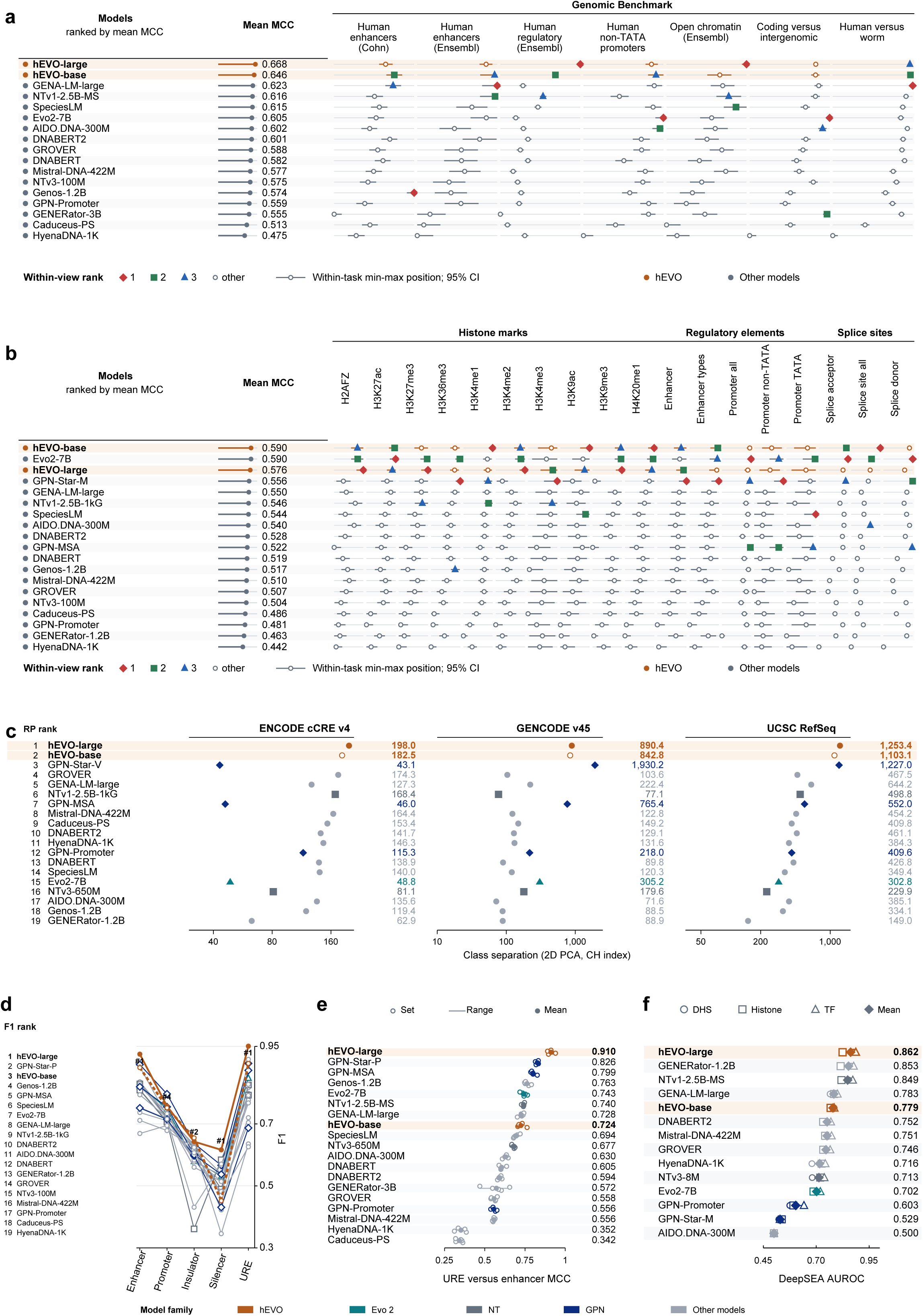
Functional genomic annotation in pretrained genomic representations. **a,b,** Frozen-encoder linear-probe performance on seven Genomic Benchmark tasks (a) and 18 Nucleotide Transformer (NT) tasks (b). Per-task positions are scaled within each task for display. Numerical summaries are arithmetic mean Matthews correlation coefficients (MCCs), with 95% test-set bootstrap intervals where available. Models lacking complete eligible-input coverage are excluded from the seven-task summary. **c,** Separation of annotation-defined classes in two-dimensional principal component analysis (PCA), quantified by the Calinski-Harabasz (CH) index. ENCODE cCRE v4 contains five regulatory annotation classes. GENCODE v45 contains coding sequences, pooled noncoding exons and protein-coding introns. Curated RefSeq contains coding sequences, 5′ and 3′ untranslated regions, protein-coding introns and noncoding transcript exons. Each collection contains 1,000 examples per class. Values summarize reduction runs with seeds 42, 43 and 44. Overall ordering uses the geometric mean of within-collection ranks. **d,** Class-specific F1 scores averaged across five regulatory datasets containing enhancers, promoters, insulators, silencers and unmarked regulatory elements (UREs). Each dataset contains 1,352 sequences per class before splitting. Ordering uses the equal-weight mean across the five classes. After an 80:20 split and removal of test sequences matching the corresponding training set under the uppercase, first-1,024-bp rule, each dataset contains 5,408 training and 1,305 test records. **e,** URE versus enhancer classification across five benchmark sets, each containing 1,355 sequences per class before splitting. Small markers show individual MCCs, larger markers their mean and connectors the observed range, not a confidence interval. The same sequence-comparison rule leaves 2,168 training and 495 test records per set. **f,** DeepSEA prediction after supervised low-rank adaptation. Markers show category AUROC for accessibility, histone modifications and transcription factor binding. The summary gives equal weight to the three categories, which together contain 919 targets. Native-input coverage and adaptation settings differ for some models. Main comparisons retain the best eligible configuration of each algorithm or version by the panel metric, together with both full-resource hEVO models.

### Haplotype learning supports broad genomic annotation and fine-grained regulatory element discrimination

Functional annotation provided a first test of whether human haplotype learning captures distinctions that extend across genomic contexts. With each encoder held fixed, linear probes evaluated seven Genomic Benchmark tasks (25) and 18 NT tasks (11), spanning regulatory elements, chromatin annotations and splice sites. hEVO-large and hEVO-base ranked first and second across Genomic Benchmark, with mean Matthews correlation coefficients (MCCs) of 0.668 and 0.646, compared with 0.623 for GENA-LM-large (26) (Fig. 2a and Supplementary Table S4). Across NT tasks, hEVO-base achieved 0.5904, closely matching Evo2-7B at 0.5896, while hEVO-large achieved 0.5758 (Fig. 2b and Supplementary Table S5). On the five shared human regulatory tasks covering enhancers, promoters, regulatory elements and open chromatin, hEVO-large and GPN-Star-M were closely matched at 0.579 and 0.578 (Supplementary Fig. S3). Without fitting a classifier, hEVO-large also showed the greatest separation of ENCODE cCRE and RefSeq classes in two-dimensional principal component analysis (PCA), with GPN-Star models performing better on GENCODE classes (Fig. 2c, Supplementary Fig. S4 and Supplementary Table S31). The strong GENCODE but weaker ENCODE separation of GPN-Star is consistent with different information being useful for gene structure and regulatory annotations, although these projections do not establish its cause. hEVO retained informative distinctions across both annotation types.

Recognizing regulatory DNA requires distinguishing elements with different roles in transcription, including promoters, enhancers, insulators and silencers. Joint evaluation of these classes tests distinctions beyond the individual promoter or enhancer tasks in standard sequence benchmarks. We also included UREs, an operational class of candidate intervals supported by genetic, functional or evolutionary evidence but lying outside selected conventional annotations (27, 28). Across five datasets containing promoters, enhancers, insulators, silencers and UREs, hEVO-large achieved the highest classification performance in every set. Its mean F1 score, which balances precision and recall with equal weight for each class, was 0.778, compared with 0.757 for GPN-Star-P and 0.725 for Genos-1.2B (Fig. 2d, Supplementary Fig. S5a and Supplementary Table S6). Recognition was strongest for UREs and enhancers, with mean F1 scores of 0.951 and 0.925. Their direct comparison tested whether the representation could distinguish candidate regulatory DNA outside conventional annotations from annotated enhancer-like elements, rather than merely separating regulatory from inactive sequence. hEVO-large achieved a mean MCC of 0.910 across five sets, compared with 0.826 for GPN-Star-P and 0.763 for Genos-1.2B (Fig. 2e, Supplementary Fig. S5b and Supplementary Table S7). These results show that the representation distinguishes multiple regulatory classes, including candidates outside conventional annotations.

Regulatory function also depends on the chromatin state and transcription factor occupancy of a sequence across cellular settings. Using DeepSEA (29), we adapted the pretrained models through low-rank adaptation (LoRA) (30) to predict 919 chromatin features. hEVO-large achieved the highest category area under the receiver operating characteristic curve (AUROC) for chromatin accessibility, histone modifications and transcription factor binding, at 0.876, 0.824 and 0.887, respectively (Fig. 2f and Supplementary Table S8). The equal-weight category mean was 0.862, compared with 0.853 for GENERator-1.2B (31) and 0.849 for NTv1-2.5B-MS. Precision-recall evaluation supported the same aggregate ordering, with mean category areas under the precision-recall curve (AUPRCs) of 0.321, 0.296 and 0.281, respectively (Supplementary Fig. S6). hEVO thus combined functional information accessible through a fixed encoder with the capacity to learn more detailed chromatin predictions after supervised adaptation.

### Variant prediction extends beyond pathogenicity to trait-associated and regulatory effects

Variant interpretation encompasses clinical pathogenicity, contribution to complex traits/diseases and changes in molecular activity, none of which is a universal measure of evolutionary fitness (32, 33, 34). We designed six complementary benchmarks to test how well pretrained scores transfer across this range without learning the evaluated outcome labels (Fig. 3a,b). They comprised ClinVar classifications (35), confirmed *de novo* loss-of-function or splice variants from the Deciphering Developmental Disorders (DDD) study (36), broader and rare fine-mapped disease-associated GWAS candidates from CAUSALdb2 (37), UK Biobank rare-variant associations with diseases and quantitative traits (19), and GWAS or expression quantitative trait locus (eQTL) candidates with regulatory effects measured by massively parallel reporter assays (MPRAs) (5). Positive and control sets were assembled from each source with allele validation, exclusion of exact class overlaps and frequency or genomic constraints where applicable (Supplementary Table S9). The same hEVO allele-likelihood contrast was used throughout, oriented towards disruption for clinical and association tasks and evaluated by magnitude for regulatory effects. This design tests generalization across distinct kinds of variant evidence rather than repeated recovery of one definition of pathogenicity.

**Figure 3.**
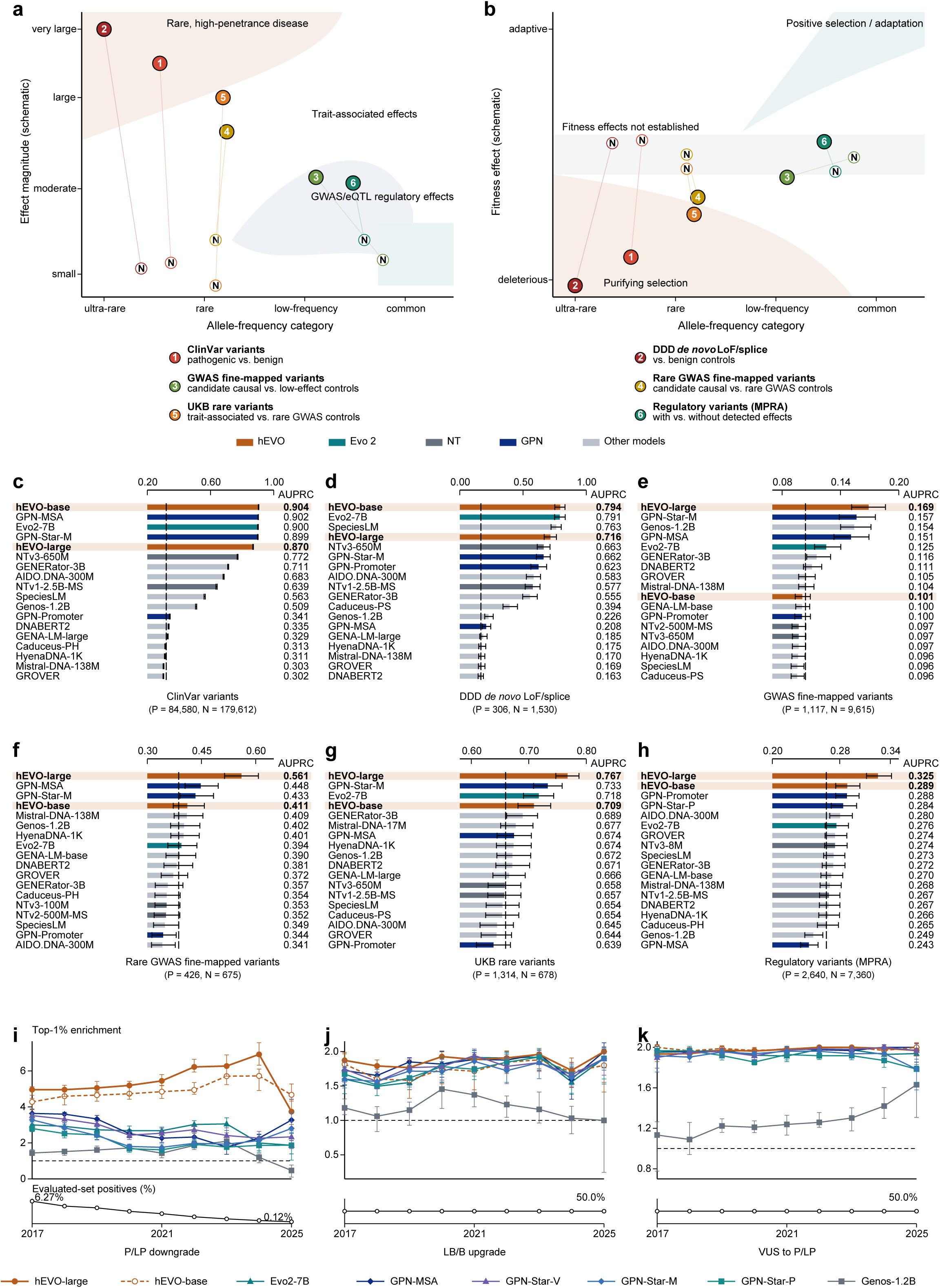
Variant prediction across clinical, association and experimental evidence. **a,b,** Conceptual relationships of the six benchmarks to phenotypic effect (a), evolutionary fitness (b) and allele-frequency categories. Numbers identify positive and control classes. Symbol positions are schematic and do not measure effect size or establish exact frequency matching. **c-h,** Zero-shot discrimination of ClinVar pathogenic versus benign variants (c), DDD *de novo* loss-of-function or splice variants versus benign ClinVar controls (d), fine-mapped GWAS candidates versus low-evidence controls (e), rare GWAS candidates versus rare GWAS controls (f), UK Biobank rare-association variants versus rare GWAS controls (g), and MPRA-supported regulatory variants versus source controls without emVar support (h). AUPRC is calculated as average precision. Whiskers indicate 95% percentile test-set bootstrap intervals where available. Positive/negative input counts for the six comparisons are 84,580/179,612, 306/1,530, 1,117/9,615, 426/675, 1,314/678 and 2,640/7,360, respectively. The corresponding input positive fractions are 0.320, 0.167, 0.104, 0.387, 0.660 and 0.264. Finite valid-score counts and average-precision baselines can differ by model. **i-k,** Retrospective top-1% enrichment for pathogenic or likely pathogenic (P/LP) variants later downgraded (i), likely benign or benign (LB/B) variants later upgraded (j), and variants of uncertain significance (VUS) later resolved as P/LP rather than LB/B (k), at annual cutoffs from 2017 to 2025. Points are means across five sampled evaluation sets. Error bars are 2.5th-97.5th percentile intervals from a two-level bootstrap that resampled sets and then variants within sets, using 20 resamples. Enrichment is the event fraction among the top-ranked 1% divided by the event fraction in the evaluated set. Dashed lines mark onefold enrichment. Lower strips show evaluated-set event fractions, not clinical prevalence.

We emphasized AUPRC because it measures the recovery of positives while accounting for false positives in these unequal-sized classes, with AUROC providing a complementary assessment (Supplementary Fig. S7). hEVO-base achieved a ClinVar AUPRC of 0.904, close to GPN-MSA at 0.902, Evo2-7B at 0.900 and GPN-Star-M at 0.899 (Fig. 3c and Supplementary Table S10). In the exploratory DDD comparison, hEVO-base achieved an AUPRC of 0.794, compared with 0.791 for Evo2-7B (Fig. 3d and Supplementary Table S11). Association-based evaluations tested a different capability, spanning a broader allele-frequency distribution and two settings focused on rare variation. hEVO-large achieved the highest AUPRC in all three, at 0.169 for broader fine-mapped GWAS candidates, 0.561 for rare candidates and 0.767 for UK Biobank rare-variant associations, compared with 0.157 for GPN-Star-M, 0.448 for GPN-MSA and 0.733 for GPN-Star-M, respectively (Fig. 3e-g and Supplementary Tables S12-S14). Its advantage extended to the MPRA benchmark, which tests measured allelic regulatory effects among GWAS and eQTL candidates, with an AUPRC of 0.325 compared with 0.288 for GPN-Promoter (Fig. 3h and Supplementary Table S15). Applying the published GPN-Star mutation-rate calibration (10) raised GPN-Star-M to 0.167 in broader GWAS and GPN-Star-P to 0.293 in MPRA, narrowing but not reversing these comparisons (Supplementary Table S29). Taken together, hEVO-base or hEVO-large achieved the highest AUPRC across all six benchmarks, supporting zero-shot transfer across clinical, association and experimental evidence rather than a single definition of pathogenicity.

Clinical interpretation also requires revisiting earlier classifications as evidence accumulates (38, 39). Across annual ClinVar cutoffs from 2017 to 2025, we evaluated subsequent downgrades of pathogenic or likely pathogenic (P/LP) variants, upgrades of likely benign or benign (LB/B) variants and resolution of variants of uncertain significance (VUS) towards pathogenicity (Fig. 3i-k and Supplementary Table S16). The largest difference concerned P/LP variants later downgraded. Among the top 1% prioritized for this outcome, hEVO-large achieved 4.96 to 6.92-fold enrichment across the 2017-2024 cutoffs. At the 2024 cutoff, enrichment reached 6.92, compared with 2.26 for the strongest external comparators, GPN-MSA and GPN-Star-V. hEVO-base performed best at the 2025 cutoff, with 4.67-fold enrichment, while the other two outcomes showed smaller differences among hEVO, Evo2-7B and the GPN models. Underrepresentation of some ancestries has contributed to erroneous pathogenic assignments (40), and later population evidence can improve classification accuracy (41). Learning naturally occurring haplotypes may help identify sequence backgrounds compatible with human variation, offering a plausible explanation for the downgrade results without implying that the underlying alleles changed their biological effects. Together, these evaluations extend the utility of hEVO from prioritizing candidate functional variants to reassessing existing clinical interpretations.

### Human sequence diversity informs population structure and lineage-specific evolution

Functional and clinical prediction do not establish whether a model retains the population differences and lineage-specific changes that shaped human genomes. We therefore extended evaluation to these less routinely tested uses of genomic language models. Ancestry analysis asks whether representations preserve variation among individuals, rather than only features shared across the species. In 1,500 individuals spanning five 1000G superpopulations, hEVO-large achieved a classification MCC of 0.985, compared with 0.983 for Evo2-7B and NTv3-650M, alongside the strongest separation in the two-dimensional t-SNE analysis (Fig. 4a, Supplementary Fig. S8a,b and Supplementary Tables S17 and S18). Continuous ancestry-proportion prediction tested information beyond group membership. With 60 training individuals per population, hEVO-large achieved a mean Pearson correlation of 0.952, compared with 0.947 for Evo2-7B and 0.944 for NTv1-2.5B-1kG (Fig. 4b, Supplementary Fig. S8c and Supplementary Table S19). These results show that sequence learning retained individual-level population information accessible with limited supervised training, complementing its functional predictions.

**Figure 4.**
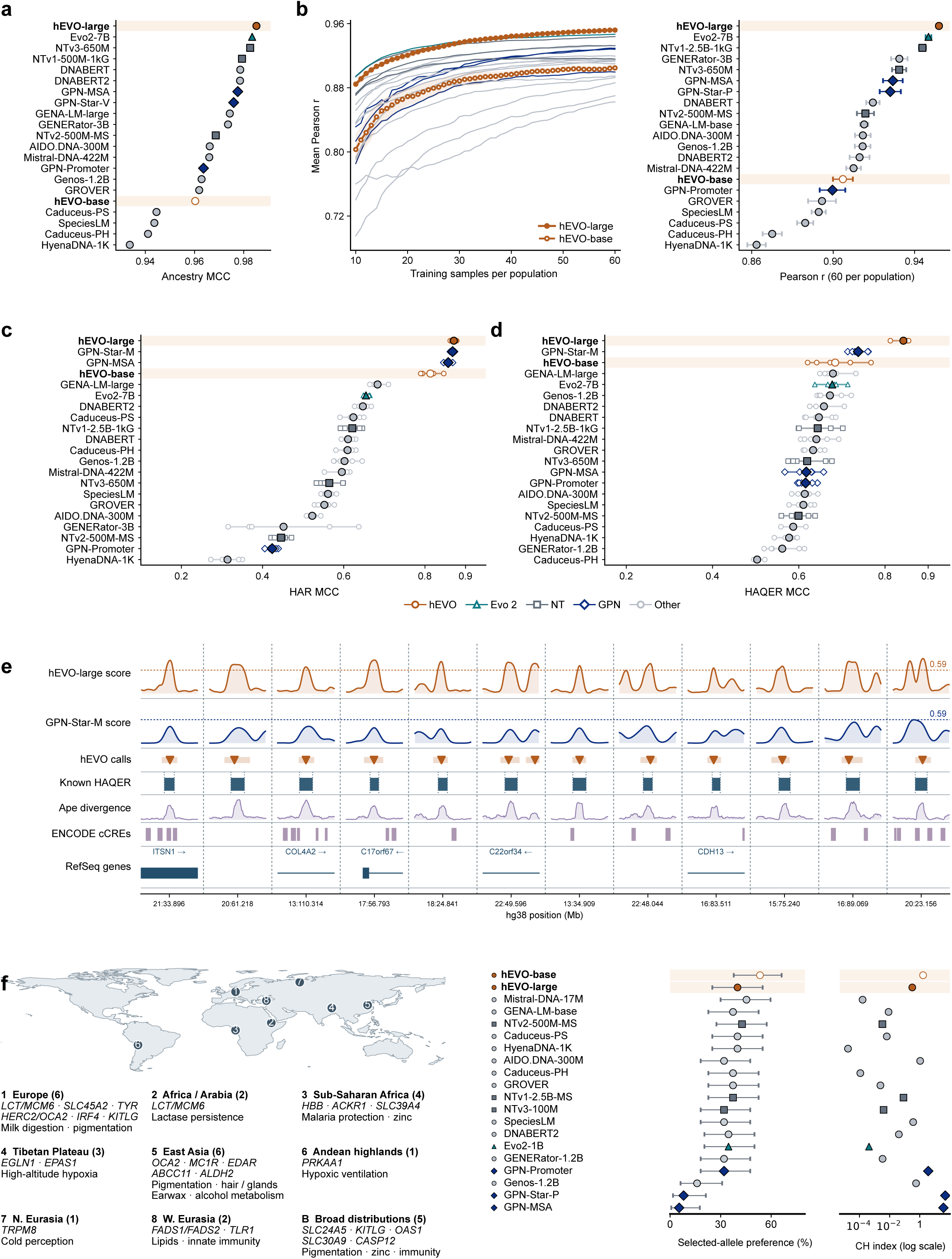
Population information and human evolutionary tracking in genomic representations. **a,** Five-group ancestry classification for 1,500 1000G individuals, with 300 from each superpopulation. MCC summarizes stratified five-fold evaluation, with feature standardization fitted within each training fold. Aggregate estimates are shown without fold-level intervals. **b,** Few-shot ridge prediction of ADMIXTURE-derived ancestry proportions, with row-wise softmax applied to the five predicted components. Learning curves show mean Pearson correlation as training size increases from 10 to 60 individuals per population. The comparison at 60 individuals per population shows means and SD across sampling repetitions, not confidence intervals. **c,d,** Linear-probe classification of human accelerated regions (HARs, c) and human ancestor quickly evolved regions (HAQERs, d) against intergenic controls. Each replicate contains 4,284 HAR-positive or 1,541 HAQER-positive sequences and equally many intergenic controls. Five replicates vary control sampling and the window-level 80:20 training/test split. The 1,541 HAQER-positive windows derive from 1,417 source intervals. Grouping by source interval and sequence deduplication were not used for this split. Small markers show individual MCCs, larger markers their mean and connectors the observed range. These are not independent pretraining runs or confidence intervals. Both classification datasets use 512 bp sequence contexts. **e,** Twelve selected, discontinuous 4 kb contexts around published HAQER intervals. Tracks show hEVO-large and GPN-Star-M scanning scores, hEVO calls, known HAQER intervals, ape divergence, ENCODE cCREs and RefSeq genes. Coordinates are hg38 positions in megabases. The validation and test contexts illustrate published HAQER intervals with qualifying hEVO calls but no GPN-Star-M calls. The first two contexts are test-set HAQER1262 in the *ITSN1* region and intergenic HAQER0258, with published comparative-chromatin and fetal-brain DNase-hypersensitivity annotations, respectively. Neither was directly tested in the original HAQER reporter assays. The threshold of 0.59 was selected using validation data. **f,** Geographic, population and functional annotations for 30 literature-informed adaptation loci, with selected allele preference and CH score separation. Preference is the fraction of loci where the reported selected allele scores above its alternative. Ties are not concordant. Intervals are exact two-sided 95% Clopper-Pearson binomial intervals. CH measures separation irrespective of direction. Colours and shapes identify model families.

Human accelerated regions (HARs) (1) and human ancestor quickly evolved regions (HAQERs) (42) provide a distinct test of human-lineage sequence change. HAQERs are especially informative because they include regulatory elements derived from ancestrally unconstrained sequence, for which deep conservation is an incomplete guide to function. hEVO-large and GPN-Star-M performed similarly in HAR classification, with mean MCCs of 0.871 and 0.868. The difference was larger for HAQERs, at 0.843 and 0.738, respectively, suggesting that haplotype-derived representations retain features useful for recognizing recently evolved sequence (Fig. 4c,d, Supplementary Fig. S9a,b and Supplementary Tables S20 and S21). In a separate chromosome-held-out scanning analysis, hEVO recovered HAQER1262 in the *ITSN1* region, annotated with human-lineage gains in regulatory chromatin during corticogenesis (42, 43), and intergenic HAQER0258, annotated with fetal-brain DNase hypersensitivity (42, 44), where GPN-Star-M did not produce qualifying calls (Fig. 4e). These independently catalogued intervals illustrate sensitivity to human-lineage regulatory contexts. Across the complete shared test set of 214 HAQERs, higher recall was accompanied by a precision trade-off (see details in Supplementary Information). The classification and scanning results together motivate the use of human haplotype representations to investigate sequence functions not readily inferred from ancestral conservation alone.

Recent adaptation raises a further distinction because a functional change does not determine which allele was favoured in a particular population or environment (2, 3). On a curated panel of 30 loci, both full-resource hEVO models preferred reported selected alleles more often than Evo2-7B, Genos-1.2B, GPN-MSA and the three GPN-Star models, with hEVO-base showing the strongest preference among these comparisons (Fig. 4f, Supplementary Figs. S9c and S10 and Supplementary Tables S22 and S23). Despite strong allele-score separation, GPN-MSA and GPN-Star usually preferred the non-selected allele, opposing the reported direction of selection. This pattern is consistent with human haplotype learning retaining sequence alternatives that need not maximize cross-species conservation, although the small panel does not establish the mechanism. Across population structure, accelerated sequence recognition and selected allele preference, hEVO brings within-human variation into genomic analyses usually assessed through functional annotation or comparative evolution alone.

### Diversity-aware learning supports compact and accessible genomic prediction

To examine which training choices contributed to this predictive breadth, we compared corpus composition, sequence overlap and allelic refinement at the base-model scale (Fig. 5a). The 63,809-individual resource achieved a mean NT MCC of 0.590, compared with 0.576 from 3,202 1000G individuals and 0.554 from 490,541 UK Biobank participants. Thus, the largest cohort did not produce the strongest representations under the evaluated training regimes. Learning overlapping sequence contexts improved performance in all three resources relative to nonoverlapping windows, whose mean MCCs were 0.559, 0.555 and 0.532, respectively (Fig. 5b and Supplementary Table S24). Improvements for the representative resource extended across histone, regulatory and splice-site tasks (Supplementary Fig. S11). Allelic refinement contributed a separate gain, improving ClinVar AUPRC in all six configurations and raising it from 0.471 to 0.904 for the representative corpus with overlapping windows (Fig. 5c and Supplementary Table S25). Because refinement preserved the encoder, it enhanced variant scoring without discarding the functional representation. These ablations support the combined value of diverse sequence backgrounds, overlapping context and focused learning at variable sites.

**Figure 5.**
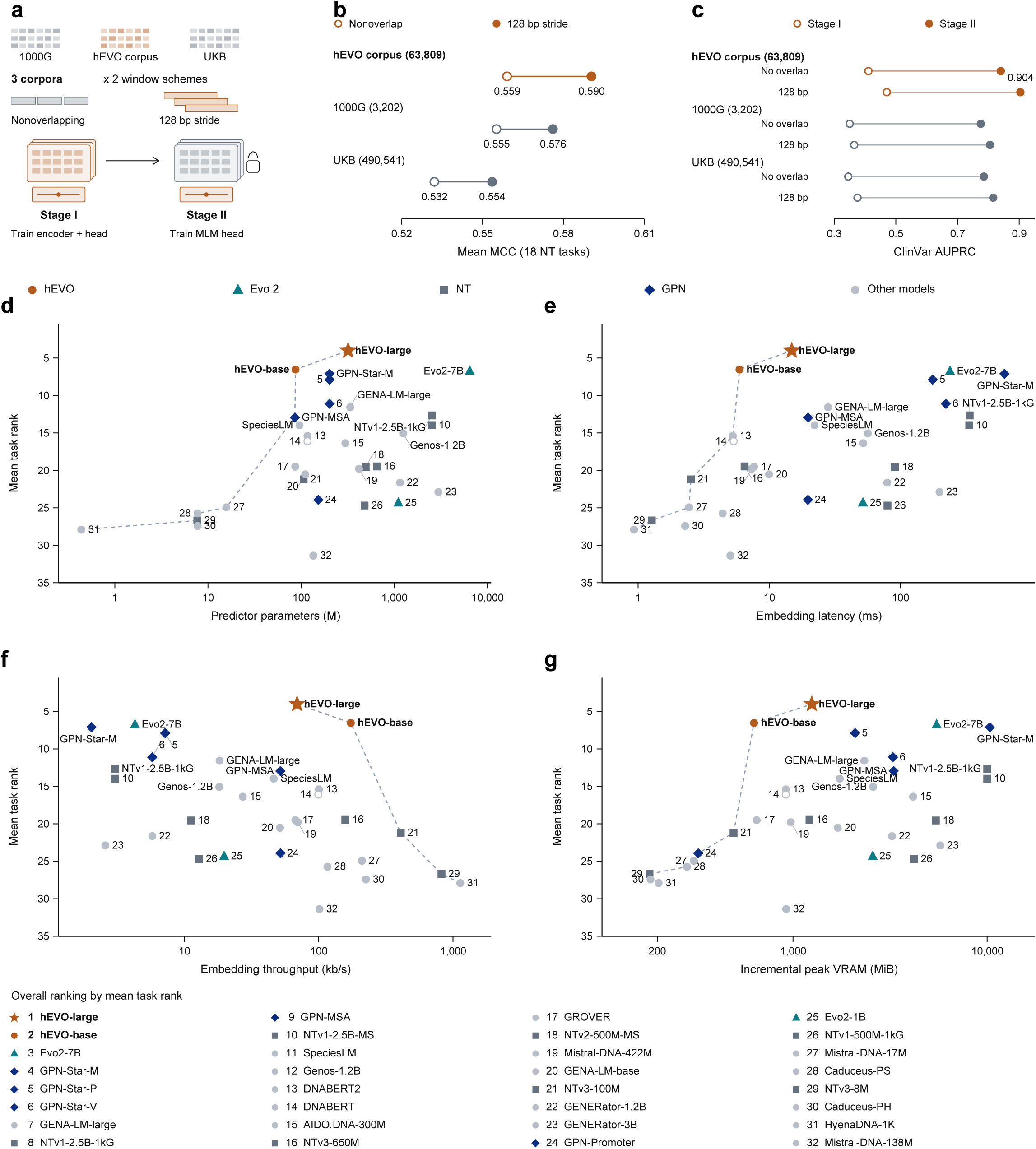
Corpus organization, allelic refinement and resource use. **a,** Six base-model ablation configurations combining three corpora with overlapping or nonoverlapping windows. The resources contain 3,202 1000G individuals, 63,809 hEVO individuals and 490,541 UK Biobank individuals. Overlapping windows are 1,024 bp long with a 128 bp stride. **b,** Mean MCC across 18 NT tasks for the six configurations. Estimates summarize task performance across the six training configurations. **c,** ClinVar average precision before and after Stage II. Lines connect the two estimates for each configuration and are not uncertainty intervals. Stage II updates the complete MLM head while preserving the sequence encoder. **d-g,** Mean task rank versus measured predictor parameters (d), embedding latency (e), nominal base-pair throughput (f) and incremental peak GPU allocation beyond model loading (g) for 32 profiled models. Mean rank uses available measurements in a prespecified 45-unit inventory, with lower values indicating better performance. The inventory includes a separate DDD same-position comparison and excludes DeepSEA, clinical reclassification and selected allele preference. Ranking units, rather than broad biological domains, receive equal weight. Resource values are arithmetic means across NT embedding workloads. Timing includes batch preparation, device transfer, embedding extraction and return to CPU. Parameter counts describe the profiled predictor and do not necessarily include the full MLM head. Memory is additional to model loading. Resource axes are logarithmic. Dashed lines connect Pareto-nondominated models with complete rank and resource coverage, not fitted trends or error bounds. Open markers denote incomplete coverage. Direct labels identify leading representatives, and the model key is ordered by mean task rank. These workload profiles are not hardware-matched speed comparisons.

Access agreements and participant consent govern the use and distribution of controlled genomic resources. To support independent deployment, we trained hEVO-base-o and hEVO-large-o using only public 1000G and HGDP data in both stages, without initialization or distillation from the full-resource models. hEVO-base-o achieved a Genomic Benchmark mean MCC of 0.627, compared with 0.623 for GENA-LM-large, and a mean regulatory-class F1 of 0.727, comparable to Genos-1.2B at 0.725. hEVO-large-o retained an NT mean MCC of 0.570 and an ancestry-proportion correlation of 0.952, although URE discrimination, association-based variant prioritization and chromatin prediction were weaker than for hEVO-large (Supplementary Fig. S12 and Supplementary Table S26). The public-resource models therefore retain useful capabilities without reproducing every benefit of the larger corpus.

The models also combined broad performance with a compact computational footprint. In the task-resolved ranking used for resource profiling, hEVO-large had the best mean rank, at 4.02 across the 45-unit inventory, and remained first when the 32 profiled models were reranked on their 38 shared units, at 3.84 compared with 6.13 for Evo2-7B and 6.39 for GPN-Star-M (Fig. 5d-g, Supplementary Fig. S13 and Supplementary Tables S27 and S28). Its measured predictor contained 319.5 million parameters and hEVO-base contained 87.1 million, approximately 20-fold and 74-fold fewer than Evo2-7B. Recorded embedding extraction averaged 14.86 and 5.93 ms per sequence, respectively, compared with 239.10 ms for Evo2-7B, with incremental peak GPU allocations of 1,251, 631 and 5,491 MiB. These measurements describe the evaluated workloads, whose device counts and execution settings differed. Other frameworks offer capabilities beyond this comparison, including the long-context and generative scope of Evo 2 (8) and Genos (12), the explicit comparative information of compact GPN-MSA and GPN-Star models (9, 10), and the broad task adaptability of NT (11). hEVO’s contribution is a combination of reusable functional representations and allelic scoring from DNA sequence alone, at a model size compatible with repeated annotation and efficient local research workflows.

## Discussion

Human genetic diversity provides evidence about genome function that complements the constraints inferred from comparisons across species. hEVO makes this evidence usable within a common sequence model, retaining distinct haplotype backgrounds and refining predictions at naturally variable sites. Leading performance across nearly all primary benchmark comparisons reflected complementary strengths of the two model scales: their representations supported functional annotation, while label-free allele scores transferred across clinical classifications, association studies and experimental regulatory measurements. Population and evolutionary analyses extended their use to differences among individuals and sequence changes within the human lineage. This breadth suggests that variation within humans can inform several forms of genomic inference, with functional recognition and variant interpretation drawing on the same pretrained representation.

Human diversity already informs NT (11), Genos (12), MutBERT (13) and UKBioBERT (14), and comparative frameworks such as GPN-Star can recover substantial complex trait information (10). Our results suggest that the organization of human variation and the objective used to learn from it also matter. Distinct-sequence sampling limits repeated exposure to common strings, while allelic refinement directs learning towards variation within those backgrounds. Naturally occurring haplotypes may consequently help represent alleles whose effects are not well described by a pathogenic-versus-neutral distinction, consistent with the separation of deleteriousness and trait modulation developed in FIND (34). The association and MPRA results support this possibility, but do not establish that hEVO estimates fitness or causal trait effects. The downgrade analysis is relevant for a related reason. Diverse population data can correct apparently pathogenic assignments (40, 41), and a model trained on such variation may supply complementary evidence for reassessment. Determining whether this explains individual hEVO predictions will require analyses that connect model scores to ancestry, allele frequency and the evidence underlying each reclassification.

The functional results point towards uses beyond broad sequence classification. Distinguishing promoters, enhancers, insulators, silencers and UREs extends annotation across regulatory classes, while chromatin prediction captures additional cellular properties. These capabilities do not yet fully specify when or where a sequence acts. Cell-specific annotations from ENCODE (45), molecular predictions from AlphaGenome (15) and functional perturbation data could help adapt population-informed representations to these contexts.

Recent personal-expression models illustrate the value of such adaptation and the continuing difficulty of transfer across loci (17, 46). Evolutionary inference presents a parallel opportunity. HAQER recognition concerns recently evolved regulatory sequence (42), whereas selected allele preference concerns the direction of change at particular loci. Strong performance on one does not ensure the other. Population-resolved training combined with historical allele-frequency evidence (2) could test which human sequence changes a model captures, provided that demographic history is distinguished from selection. The 30-locus analysis supports further evaluation of adaptive allele preference using larger, population-resolved panels.

The contrast between model sizes reinforces the need to assess these capabilities separately. hEVO-base retained stronger ClinVar discrimination and selected allele preference, whereas hEVO-large improved several regulatory and association-based tasks. The public-resource base model also matched the full-resource base model on selected allele preference. These patterns do not show that smaller models or fewer genomes are intrinsically preferable, because training exposure, optimization and refinement differed and the selection panel was small. A model that predicts local nucleotides more accurately need not better recover a particular clinical or evolutionary distinction. Evo 2 likewise reported saturation or declines in some molecular fitness predictions at its largest scales (8), and Genos showed task-dependent gains with model size (12). Controlled scaling experiments could distinguish the contribution of capacity from the information emphasized during training. In the meantime, the breadth attainable with compact hEVO models makes such experiments and downstream adaptation more accessible.

Several limitations define the scope of these findings. First, benchmark design affects what a performance difference can establish. Coding and noncoding variants were evaluated jointly for genome-wide prioritization, because important disease effects are not confined to coding sequence (47). Aggregate scores can nevertheless reflect consequence composition, while regulatory classification can reflect sequence length as well as function. Exact-string filtering in the URE tasks does not exclude all overlapping loci, and the window-level HAQER classification split does not establish locus-independent generalization. Second, held-out task labels do not exclude pretraining exposure to related sequences or individuals, so the population and retrospective analyses are not tests of unseen populations or prospective clinical utility. Third, the approximately 1 kb context limits distal regulation and extended-haplotype modelling. Population-variant learning was restricted to SNVs, while indels and more complex structural variants were not modelled because their reconstruction poses additional challenges in phasing and sequence representation. Fourth, Evo2-40B and Genos-10B were not deployed because of computational demands and evaluation time, and AlphaGenome requires comparisons tailored to its supervised molecular outputs. Conclusions therefore concern the evaluated checkpoints and tasks. Finally, workload-specific resource measurements and the absence of independent clinical validation limit deployment claims. Within this scope, hEVO provides a compact foundation for studying how human sequence diversity can improve functional annotation and variant interpretation.

## Methods

### Human genomic resources and cohort preparation

We assembled WGS data for 550,237 participants from UK Biobank (19), TOPMed (20), HGDP (18), 1000G (21), UK10K (22), SG10K (23), CAAPA (24) and an internal cohort of 3,188 healthy Han Chinese individuals sequenced at 30x depth. The collection retained the population-based and disease-focused ascertainment of its source cohorts and was not restricted to healthy individuals. Selection using UK Biobank ancestry codes yielded 25,934 participants enriched for non-European backgrounds; eligible individuals from the other cohorts were retained after quality control and duplicate removal, giving 63,809 individuals.

Cohort VCFs were harmonized to GRCh38, with reference checks, multiallelic decomposition and missing-genotype filters applied before sequence reconstruction. TOPMed duplicate or twin pairs were handled separately. Genetic ancestry was annotated after selection using common-variant principal components calculated with PLINK2 (48) and a calibrated, reference-trained nearest-neighbour classifier on six components. The internal Chinese cohort was assigned to East Asian ancestry from cohort information. UK Biobank SNVs were phased with SHAPEIT5 (49), while the other cohorts supplied phased genotypes (Supplementary Table S1; see Details in Supplementary Information).

### Haplotype reconstruction and diversity estimation

Phased SNV alleles were substituted into GRCh38 to reconstruct both chromosome copies in 1,024 bp autosomal windows at 128 bp steps. Windows containing ambiguous bases or unresolved genotypes were excluded. Reference filling at absent variant-only VCF records was a reconstruction convention, not evidence of read coverage at every position. Exact sequence duplicates were merged across individuals and loci, retaining occurrence counts, ancestry annotations and genomic origins; reverse complements remained distinct. Stage I sampled distinct sequences uniformly, reducing repeated exposure while changing natural sequence abundance.

To assess the diversity represented by cohort selection, we compared local SNV haplotypes in the UK Biobank subset with a 50,000-person reference sample from the full phased cohort. Separate 100 kb windows were used for this analysis, rather than the training windows. Abundance-weighted coverage measured the fraction of reference chromosome copies represented above each frequency threshold, while set coverage weighted distinct haplotypes equally. Cohort accumulation used 577 deterministically sampled, variant-bearing windows, with equal chromosome contributions from each cohort unit and ten rarefaction draws. Total chromosome count therefore increased as cohorts were added. Corpus-wide occurrence and ancestry-sharing summaries were calculated separately from the complete deduplicated resource (Supplementary Table S2; see Details in Supplementary Information).

### Model architecture and sequence pretraining

hEVO uses a bidirectional Transformer and nucleotide-level masked language modelling (50), allowing each position to draw on sequence context on both sides. The base configuration contains 12 Transformer blocks with 768-dimensional hidden states, while the large configuration contains 16 blocks with 1,280-dimensional states, corresponding to approximately 87 and 319 million parameters. The sequence encoder feeds an existing prediction head comprising a dense transformation, GELU activation, LayerNorm and an untied vocabulary decoder.

During Stage I, 20% of valid nucleotide positions were masked in short contiguous spans, and the encoder and prediction head were trained jointly to recover the hidden bases. Cross entropy was calculated only at masked positions. Distinct haplotypes supplied the training examples; measured ancestry and allele frequency were not backbone inputs. Optimization used scale-specific settings and sequence budgets, with validation separated from the training records. Architectural dimensions, masking distributions, optimization settings and training budgets are documented in Supplementary Table S30 (see Details in Supplementary Information).

### Allelic refinement and public-resource models

Stage II focused prediction on naturally variable positions using biallelic SNVs from the phased high-coverage 1000G panel. Each example retained the surrounding sequence of an observed ALT-carrying chromosome, with the focal nucleotide masked. Equal sampling across five ALT-frequency strata prevented common variants from dominating this refinement set. Whole 1 Mb genomic blocks were assigned to training, validation and test sets in an expected 90:5:5 ratio, excluding contexts that crossed block boundaries. This isolated Stage II contexts across splits but did not remove Stage I exposure to the same regions.

We kept the sequence encoder fixed and updated the complete existing MLM prediction head, including its transformation, LayerNorm and decoder. Refinement therefore changed how the learned sequence features were translated into allele probabilities without training a second encoder. Neither stage used clinical or functional outcome labels, and frequency strata were sampling information rather than model inputs. Public-resource counterparts, hEVO-base-o and hEVO-large-o, were trained independently using only 1000G and HGDP in both stages, without initialization or distillation from restricted-resource models (Supplementary Tables S26 and S30; see Details in Supplementary Information).

### Comparator interfaces and frozen representations

The comparison inventory comprised 34 checkpoint entries, including four hEVO models and representatives of Evo 2 (8), NT (11), Genos (12), GPN-MSA (9), GPN-Star (10) and other model families. Eligibility was determined for each task. Single-sequence models used their native tokenizers, whereas alignment-dependent models received the required genomic coordinates and multispecies inputs. Models lacking compatible inputs were not included in the corresponding aggregate comparison.

Functional information acquired during pretraining was assessed with the encoder fixed. Final-layer representations were pooled over supported positions, excluding padding, and an affine linear classifier was fitted to these features. Sequence-classification probes used a shared 50-epoch protocol with batch size 32. Native context limits, tokenization and pooling policies were retained and recorded rather than forcing all architectures through one sequence interface (Supplementary Table S3; see Details in Supplementary Information).

### Functional sequence and chromatin benchmarks

Genomic Benchmark (25) contributed seven tasks and the NT benchmark (11) contributed 18 tasks spanning regulatory elements, histone annotations and splice sites, with their original partitions retained. Alignment-dependent comparisons used the same five eligible human Genomic Benchmark tasks. Fine-grained regulatory evaluation included unmarked regulatory elements (UREs), an operational class informed by candidate elements outside conventional annotations (27, 28). Candidates integrated CRISPR, MPRA, eQTL and evolutionary evidence after subtraction of selected conventional epigenomic annotations. Five binary datasets compared UREs with length-bin-sampled enhancers, and five multiclass datasets combined UREs with enhancers, promoters, insulators and silencers. URE test records whose retained input sequence matched a training sequence were excluded. Multiclass categories were balanced in count but not matched in length.

Representation structure was evaluated without fitting a classifier using three annotation collections from ENCODE (45), GENCODE (51) and RefSeq (52), each containing 1,000 examples per class in 512 bp windows. Class separation was quantified by the Calinski-Harabasz index on two-dimensional PCA coordinates; complementary projections were retained in the supplementary analyses. DeepSEA (29) tested supervised adaptation through LoRA (30) across 919 chromatin targets, retaining the source training, validation and test partitions. Target-wise AUROCs and average precision were averaged within accessibility, histone modification and transcription factor binding categories, with equal category weights in the overall summary (Supplementary Tables S4-S8, S31 and S32; see Details in Supplementary Information).

### Variant benchmarks and zero-shot allele scoring

We constructed six benchmarks to separate clinical interpretation, association evidence and measured regulatory effects. ClinVar (35) supplied pathogenic and benign SNVs, while DDD (36) supplied confirmed *de novo* SNVs with predicted loss-of-function or splice consequences, compared with benign ClinVar controls. Broader and rare GWAS benchmarks used disease-associated CAUSALdb2 candidates (37). Positive candidates required a mean stored posterior above 0.8 and absolute effect estimate above 0.5; control candidates required a posterior below 10^−10^ and absolute effect below 0.01. Controls with avgPP = −1, the missing-LD code, were excluded from the broader benchmark. Broader-set controls were sampled by chromosome and nearest available allele-frequency bin, whereas the rare comparison retained eligible controls with gnomAD frequency below 1% without frequency matching. A separate UK Biobank benchmark used published sentinel associations (19) selected at an NFE effect-allele frequency below 1%, with rare GWAS controls. MPRA positives had experimentally detected allelic regulatory effects among fine-mapped GWAS or eQTL candidates, and source-category controls lacked a detected allelic effect (5).

Variant keys, alleles and coordinates were reconciled against GRCh38, with exact positive-control overlaps removed. Paired benchmark sequences differed only at the focal nucleotide in a common reference background, unlike the carrier-haplotype backgrounds used in Stage II. No classifier was fitted to these outcome labels. Masked models contrasted conditional ALT and REF probabilities, while autoregressive models used their native sequence-likelihood contrasts; documented native scorers were retained where these differed. Scores were oriented towards predicted disruption for clinical and association tasks and towards effect magnitude for MPRA. GPN-Star mutation-context calibration (10) was compared with raw scores on identical eligible variants (Supplementary Tables S9-S15 and S29; see Details in Supplementary Information).

### Retrospective clinical reclassification

Archived ClinVar (35) releases from 2017-2025 were compared with the 15 June 2026 release after matching allele-specific variant keys. Three analyses contrasted pathogenic or likely pathogenic variants later reassigned to uncertain or benign categories with stable pathogenic classifications, benign or likely benign variants later reassigned to uncertain or pathogenic categories with stable benign classifications, and uncertain variants later resolved as pathogenic rather than benign. Conflicting interpretations were excluded, without an additional star-rating threshold. Five sampled evaluation sets were generated for each cutoff and outcome, with score direction defined separately for each transition.

The event fraction among the top-ranked 1% was divided by the event fraction in the same evaluated set, so enrichment was measured against the retrospective sampling baseline rather than clinical prevalence. Point estimates averaged the five sets; percentile intervals used two-level resampling of sets and paired variant labels and scores. Repeated annual appearances of a variant were retained (Supplementary Table S16; see Details in Supplementary Information).

### Population representations and accelerated human sequences

Population analyses included 1,500 1000G individuals, with 300 from each superpopulation. Individual representations summarized observed-minus-reference features across a shared panel of 10,000 marker contexts. Five-group classification used stratified five-fold cross-validation, with standardization fitted within each training fold. Few-shot ridge regression predicted ADMIXTURE-derived proportions (53) from 10-60 labelled individuals per superpopulation; Pearson correlations were summarized across ten sampling repetitions. Projected representation geometry was assessed separately.

HAR and HAQER (42) classification used 512 bp windows paired with balanced intergenic controls in five sampling-and-split datasets. Linear probes were trained on frozen representations after window-level splitting. The 1,541 HAQER-positive windows originated from 1,417 catalogue intervals; this classification design did not isolate source loci across partitions. A distinct HAQER scanning analysis compared hEVO-large and GPN-Star-M, using chromosomes 1-12 for training, 13-17 for validation and 18-22 for testing. Prediction heads, calibration and peak-calling settings were selected using validation data and fixed before evaluating 214 shared test intervals. Window discrimination and interval detection were reported separately; selected display contexts illustrated model discrepancies and were not the aggregate test set (Supplementary Tables S17-S21; see Details in Supplementary Information).

### Preference for reported selected alleles

Thirty literature-informed loci were curated with separate annotations for allele-level selection, functional evidence and association with a selected haplotype. We reconciled selected and alternative alleles to the GRCh38 plus strand, including cases where the selected allele was the reference base. Paired sequences shared the same surrounding context, and native strand-aggregation procedures were retained. Preference was the fraction of valid loci where the selected allele scored strictly above its alternative; ties were not concordant. Exact 95% binomial intervals and a 50% random-orientation reference accompanied these counts. Paired score tests and a direction-independent CH analysis provided complementary summaries. These evaluations assessed model preference at the curated loci, not fitness or genome-wide discovery of selection (Supplementary Tables S22 and S23; see Details in Supplementary Information).

### Training ablations and computational profiling

Corpus and windowing ablations held the base architecture fixed across three resources: 3,202 1000G individuals, the 63,809-individual hEVO corpus and 490,541 UK Biobank participants. Each resource was processed with overlapping or nonoverlapping windows, giving six configurations evaluated on the 18 NT tasks. ClinVar comparisons before and after Stage II isolated refinement within each configuration. Training exposure differed between resources, so the corpus comparison did not isolate population size alone.

Computational profiling covered embedding extraction for 32 models across the NT workloads. Recorded measurements included predictor parameters, elapsed extraction time, throughput and peak GPU allocation above the loaded-model baseline. Timing included preparation, device transfer, representation extraction and transfer back to CPU. Device count, numerical precision, batching and native context limits were recorded where available; comparisons describe these workloads rather than matched-hardware training costs (Supplementary Tables S24, S25 and S27-S28; see Details in Supplementary Information).

### Statistical summaries and reproducibility

Performance summaries used MCC, equally weighted class F1, AUROC and average precision, labelled AUPRC in the figures. Metrics retained finite valid predictions, with model-specific sample counts and positive fractions; the latter defined the average-precision baseline. Ordinary test-set intervals used 1,000 paired bootstrap resamples. Control-and-split datasets, ancestry folds, repeated training-sample draws and curated selection loci had distinct sampling units and uncertainty summaries.

The headline assessment counted 19 primary comparisons: eight functional, six variant and five population or evolutionary endpoints. A comparison was led by hEVO when either full-resource model had the highest eligible point estimate. Resource analyses separately averaged ranks across 45 available units in the 34-model inventory, with a sensitivity analysis recomputing 38 shared units among 32 profiled models. Radar summaries retained 13 axes and selected the best eligible family member on each axis, with all GPN algorithms contributing to the GPN family. Main panels retained both full-resource hEVO models and eligible algorithm representatives. Dataset versions, checkpoints, input definitions, score directions and seeds were recorded for reproducibility; access and release arrangements are specified under Data and code availability (see Details in Supplementary Information).

## Acknowledgements

The work was supported by the National Key Research and Development Program of China (2024YFC2707700 to M.J.L., and W.H.Z.) and the National Natural Science Foundation of China (32470671 to M.J.L.). We thank the participants and investigators who contributed to the UK Biobank, the NHLBI Trans-Omics for Precision Medicine (TOPMed) program, the UK10K project and its contributing ALSPAC and TwinsUK cohorts, the SG10K_Pilot study, and the Consortium on Asthma among African-ancestry Populations in the Americas (CAAPA). We acknowledge the contributions of the 1000 Genomes Project and the Human Genome Diversity Project to publicly available genomic reference resources, and thank the participants and colleagues who contributed to the internal Chinese cohort. We acknowledge the National Heart, Lung, and Blood Institute for supporting TOPMed, and the TOPMed Informatics Research Center and Data Coordinating Center for their roles in genomic data processing and program coordination. We also acknowledge the core support for ALSPAC from the UK Medical Research Council, Wellcome, and the University of Bristol, and the support for TwinsUK from the Wellcome Trust, Medical Research Council, Versus Arthritis, the European Union Horizon 2020 programme, the Chronic Disease Research Foundation, the Wellcome Leap Dynamic Resilience Programme, Zoe Ltd, and the National Institute for Health and Care Research Clinical Research Network and Biomedical Research Centre at Guy’s and St Thomas’ NHS Foundation Trust in partnership with King’s College London. The SG10K_Pilot data were obtained from EGA; this work was not prepared in collaboration with the SG10K_Pilot study and does not necessarily reflect its views.

## Author contributions

M.J.L. conceived the study. Q.X., Z.W., and M.D. processed the data, developed the model, performed benchmarking, and drafted the manuscript. W.N., X.H., H.Y., Y.Z., and X.Y. contributed to data access and processing. T.B., and S.Z. contributed to model development, and D.H. performed variant analyses. P.C.S. provided scientific guidance. W.Z. and M.J.L. supervised the study and contributed to writing and revising the manuscript.

## Data and code availability

Code and analysis workflows are available at https://github.com/mulinlab/hEVO. Numerical benchmark results and source definitions accompany the article. Participant-level genomes remain subject to cohort agreements, consent and approved research purposes; repository licences and upstream data conditions govern reuse. Non commercial.

